# Replicated point processes with application to population dynamics models

**DOI:** 10.1101/409292

**Authors:** Marco Favretti

## Abstract

In this paper we study spatially clustered distribution of individuals using point process theory. In particular we discuss the spatially explicit model of population dynamics of Shimatani (2010) which extend previous works on Malécot theory of isolation by distance. We reformulate Shimatani model of replicated Neyman-Scott process to allow for a general dispersal kernel function and we show that the random immigration hypothesis can be substituted by the long dispersal distance property of the kernel. Moreover, the extended framework presented here is fit to handle spatially explicit statistical estimators of genetic variability like Moran autocorrelation index, Sørensen similarity index, average kinship coefficient. We discuss the pivotal role of the choice of dispersal kernel for the above estimators in a toy model of dynamic population genetics theory.

## 1. Introduction

Spatial point processes [1, 2] are a valuable tool for quantitative analysis and model design in population genetics and community ecology studies. They are useful for discussing central issues such as isolation by distance, clustering, distance decay of similarity and more generally in all cases when the spatial distribution of individuals and spatial genetic variability matters. In this paper we focus on the specific class of spatial processes called Neyman-Scott processes [20], see Section 2 below. In these, each point of a given random distribution (called parent points) generates offspring points that are dispersed away from the parent location with a prescribed probability distribution, called dispersal kernel. In a *replicated* Neyman-Scott process, offspring become new parent points while the previous generation ones are discarded. In this way we build a spatially explicit dynamic genetic model of non-overlapping generations clustered around parent points. Of course the choice of the dispersal kernel function is a crucial issue to get a realistic description of the spatial pattern and in effect there is a large literature dealing with the theoretical choice and experimental validation of this key tool (see [13] and the review [12]).

The original idea behind this model can be traced back to Malécot theory of isolation by distance [4, 5, 6]. Felsenstein [3] criticized the use of two-dimensional Gaussian probability density as the dispersal kernel. This choice seems the most natural one in view of the nature of the computation but it produces an unbounded clustering. Therefore the model was rejected by Felsenstein as physically unsound (see however the comment in [7] to Felsenstein paper). Recently the model has been reformulated by Shimatani [8] in the spatial point process framework still considering a Gaussian dispersal kernel but adding the hypothesis that at each generation a fixed fraction of offsprings migrates randomly away from their parents points. In this way the resulting process reaches a finite clustering even if an infinite number of generations is considered.

In this paper we repeat the computation along the lines of [8] for a general dispersal kernel function. We thus obtain a model of greater generality in which we are able to discuss the role of the choice of dispersal kernel on the issue of divergent clustering. We can show that the use of a leptokurtic dispersal kernel (i.e. with a decay slower than the gaussian or exponential function) do produce a finite clustering without the need of the random migration hypothesis, which is thus replaced by a *long distance dispersal* hypothesis. Gaussian and exponential kernels were commonly considered until the turning of the century, but nowadays are outperformed by various fat tailed dispersal kernels (see Ch.15 in [12]).

Replicated point process as pioneered by Shimatani [8, 9, 10] allow to consider a dynamic and spatially explicit population genetics model. In this way a coalescent theory is recovered. A powerful feature of replicated point process is that they allow to study distance-related statistical properties of spatial *and* genetic patterns. If the population is described by a marked point process (*x, m*(*x*)) where the qualitative or quantitative mark *m*(*x*) encodes the genetic information of the individual sitting at *x*, genetic variability on a given spatial pattern can be measured by the so called spatial correlation Moran Index [17] while the degree of relatedness between a couple of offsprings can be measured by their average kinship coefficient *G*, see Section 3 below. In Section 3 we consider a replicated Neyman-Scott process where genetic variability is described by a qualitative mark which represents the genotype of the individual. We assume that the probability that a parent of genotype *i* produces an offspring of genotype *j* is given by a Markov matrix *Q*_*ij*_ and that the vector of genotype frequencies *q* is a stationary distribution for *Q*. In this way genotype frequencies are constant from one generation to the other. We compute the pair correlation function between offsprings of genotype (*i, j*) for each generation. As a measure of genetic variability we compute for our model the point process formulation of Moran spatial autocorrelation index (see [9]) and Sørensen similarity index and their infinite generation limit in the case of an ergodic Markov chain. In a specific example where the qualitative marks represents the allele at a single locus, we compute explicitly the Moran index for three different dispersal kernels (Gaussian, exponential, Cauchy) and we show that a long distance dispersal kernel has a long range of spatial genetic correlation.

## 2. Replicated Neyman-Scott process with a general dispersal kernel

We briefly recall the basic facts of Neyman-Scott point processes (see and [2] for an introduction). They are stationary and isotropic point processes in the plane defined by the following steps

a) parents points are distributed on the plane with an homogeneous Poisson point process of intensity *λ*, therefore they represent a collection of points distributed at random uniformly and independently
b) each parent points produces a number *u* of daughters sampled from a given probability distribution (usually a Poisson distribution) of given expected value *U* = *E*(*u*) and variance *V.*
c) daughter seeds are dispersed from parents location using a two-dimensional radially symmetric probability *h*(*r*) (the so called *dispersal kernel*). The probability that a daughter of a parent located at the origin *O* lands on an infinitesimal area *dx* centered around point *x* in the plane is

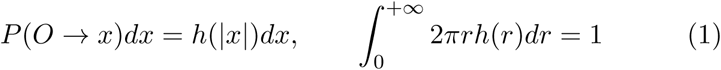

Parent point are removed, therefore the intensity (i.e expected spatial density of individuals) of the first generation is *λU.*

In a replicated Neyman-Scott process, the first generation daughters become parent points and the steps *b*) and *c*) are repeated *n* times to form the spatial distribution of individuals after *n* non overlapping generations. Note that only the initial *t* = 0 generation points are randomly distributed whereas the subsequent *t* = 1,*…, n* generations points forms clusters of individuals. A statistical description of the spatial distribution of individuals can be given along the following lines. Let *n*(*x*) and *n*(*y*) be the random variables representing the number of individuals (points) falling in infinitesimal disjoint circular regions *dx* and *dy* around points *x* and *y*. Let *λ* be the average spatial density of points of the process (first order intensity) and *ρ*(*x, y*) the so called second order product density. The following interpretations are standard (hereafter *P* (*x*) denotes probability density with respect to the area measure *dx*)

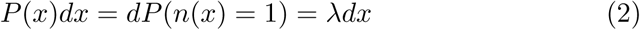

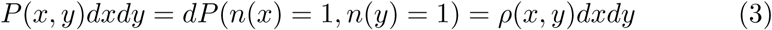

The pair correlation function *g* measures the ratio of the probability of finding two individuals at *x* and *y* in a clustered distribution and in a completely random one, therefore it gives a measure of the degree of spatial clustering

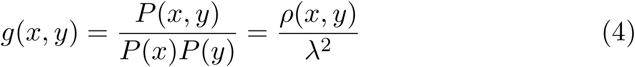

For the *t* = 0 generation, which is described by an homogeneous isotropic Poisson process we have *g* ≡ 1, while for the subsequent ones we may have *g >* 1 but the distribution is still stationary and isotropic (i.e rotation and translation invariant), therefore we may always assume that *P* (*x*) = *P* (*|x|*) and *P* (*x, y*) = *P* (*y − x*) = *P* (*|y − x|*) throughout. By requirement *c*) above, we have that after *t* generations, the expected density of individuals is (see (2) and *c*) above)

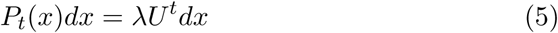

Our main aim is to compute the joint presence probability *P*_*t*_(*x, y*) for *t* ≥ 1. The following computation can be considered the generalization of the one contained in Shimatani paper [8] which is the primary source of inspiration for this work. The technical part of the computation is contained in the Supplementary material, Sect. 5.

Let *x*_1_ and *x*_2_ be the location points of two daughters at the *t* generation. They may be offsprings of different mothers (dm) located at *x* and *y* or be offsprings of the same mother (sm) at *x*. These are exhaustive and mutually incompatible alternatives, therefore we can write

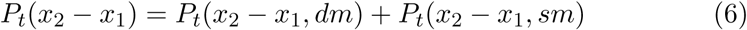

For the (*sm*) case we have, writing *P*_*t-*1_(*x*) = *λU*^*t−*1^, see (5),

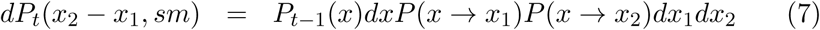

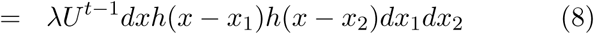

to be integrated in *dx* over the plane and multiplied for the average number of couple of daughters produced at *x E*(*u*(*u* – 1)) = *U*^2^*W* where *W* = (*U*^2^ – *U* + *V*)*/U*^2^ to give

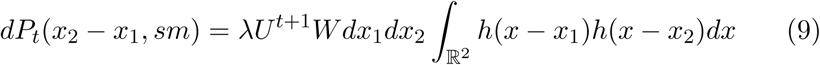

By writing *x – x*_2_ = *x*_2_ – *x*_1_ *-* (*x – x*_1_) and *dx* = *d*(*x – x*_1_) the above integral represents the convolution of *h* with itself

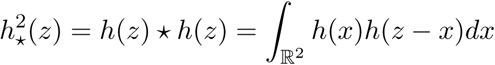

For the sake of simplicity, we consider the *dx*_1_*dx*_2_ density of the above function (9)

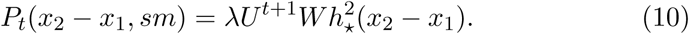

For the (*dm*) case we have

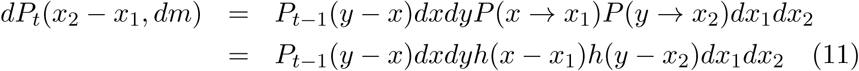

to be integrated in the area measure *dxdy* over all the possible parent locations and multiplied for the average number of daughters produced by a parent at *x* and *y*, i.e. *E*(*u*)^2^ = *U*^2^. Passing to the *dx*_1_*dx*_2_ density as above, we have

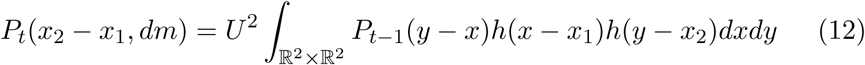

While for the (*sm*) case we have obtained the closed formula (10), for the (*dm*) case the probability *P*_*t*−1_(*y – x*) in (12) has to be computed recursively from (6) and (10). Let us do the computation for a few generations. Since the initial distribution of individuals is random *P*_0_(*y – x*) = *ρ*(*x, y*) = *λ*^2^ we have easily (using Fubini’ Theorem and (1)) that for the *t* = 1 generation *P*_1_(*x*_2_ – *x*_1_, *dm*) = *λ*^2^*U*^2^ hence from (6)

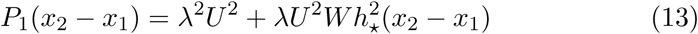

For the sake of simplicity, from now on we set *r* = *x*_2_ – *x*_1_. For the *t* = 2 generation, we have from (10) 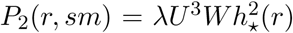 and, inserting (13) in (12) we get

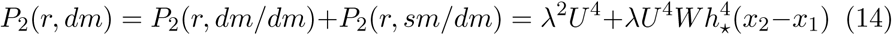

For the *t* = 1 generation we had two alternatives: (*sm*) corresponding to half-siblings, and (*dm*) corresponding to independent lineages; for the *t* = 2 generation we have three alternatives: (*sm*), (*sm/dm*) corresponding to first cousins with a common grandmother or (*dm/dm*) for independent lineages. By iterating the same computation (see Supplementary Material) we find that at generation *t* we have the following *t* + 1 exhaustive and mutually incompatible alternatives:

i) (*dm*)^*t*^ corresponding to independent lineages (no common ancestor)

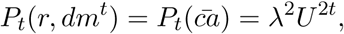
ii) *sm/*(*dm*)^*k*^ corresponding to a common ancestor *k* generations before for *k* = 1,*… t*, with probability

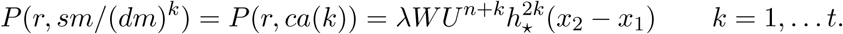

Therefore, the probability of having a common ancestor (*ca*) for two individuals of the generation *t* at distance *r* is

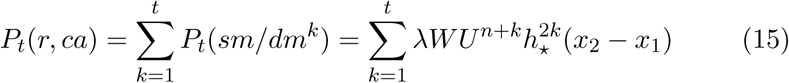

while the probability that they represent independent lineages 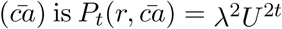. Their sum gives the joint probability of finding two individuals at distance *r*

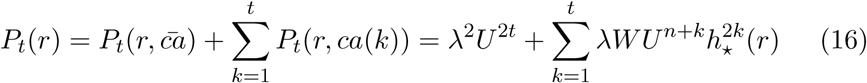

If we divide the above equality by *P*_*t*_(*r*), using the well known relation *P*(*B|A*) = *P* (*A, B*)*/P* (*B*)

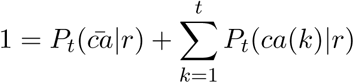

we can compute the conditional probability that two individuals belongs to different lineages given that they are at distance *r*

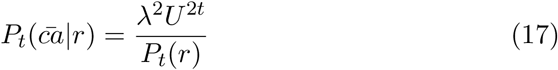

or the conditional probability that they share a common ancestor *k* generations ago given that they are at distance *r*

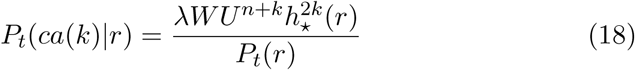

All these formulas are useful for spatial genetic applications and they show that the replicated Neyman-Scott process model is an instance of a spatially explicit coalescent theory, as pointed out by [8].

### 2.1. The long distance dispersal hypothesis

Here we expose the main contribution of this paper. The *t* generation pair correlation function computed from its definition (4) is

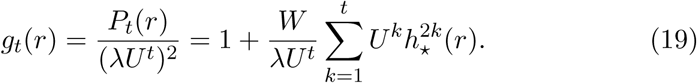

We see that the behavior of the pair correlation function (19) depends on the convergence of the series of function with positive terms 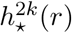.

Before considering the possible choices of the dispersal kernel *h*, let us note that in this model the spatial density of individuals after *t* generations is *λU*^*t*^, therefore in absence of any regulatory mechanism the density diverges if the fecundity is *U >* 1 while the population goes extinct if *U <* 1. In the forests the density of plants is nearly constant, hence it is reasonable to assume *U* = 1 throughout in our model. In this case *W* coincides with the variance *V* and the pair correlation function depends on the ratio *V/λ*

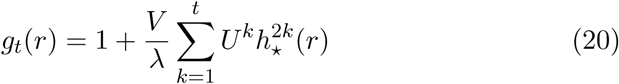

In the previously cited papers, the dispersal kernel was assumed to be a 2-dimensional Gaussian probability density

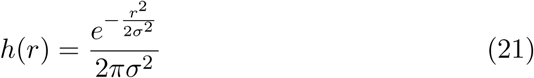

whose 2*k*-fold convolution as it is well known is still a Gaussian with variance *kσ*^2^ that is

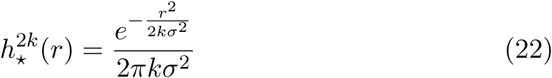

It is easy to see that, in the limit *t* → ∞ the series of function in (19) diverges since for every *r* ≥ 0

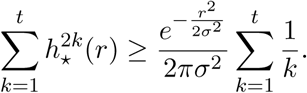

Therefore, in absence of other mechanisms, the degree of clustering diverges for *t* → ∞. See however Figure 1 below: after 1000 generations the value of the pcf at *r* = 0 is only 2.6 times the value at *t* = 0, therefore the divergence phenomena could not be visible for realistic values of the parameters.

**Figure 1:**
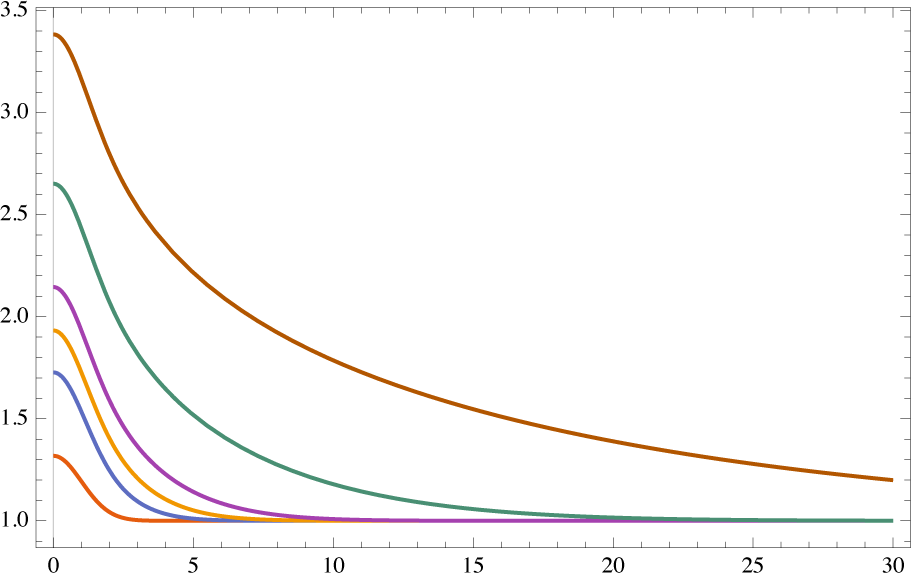
Pair correlation function *g*_*t*_(*r*) for a Gaussian kernel with parameters *σ* = 1, *λ* = 0.05, *V* = 0.1, (*V/λ* = 2) computed for *t* = 1, 5, 10, 20, 100, 1000 generations. The pcf grows unbounded with the number of generations.

**Figure 2:**
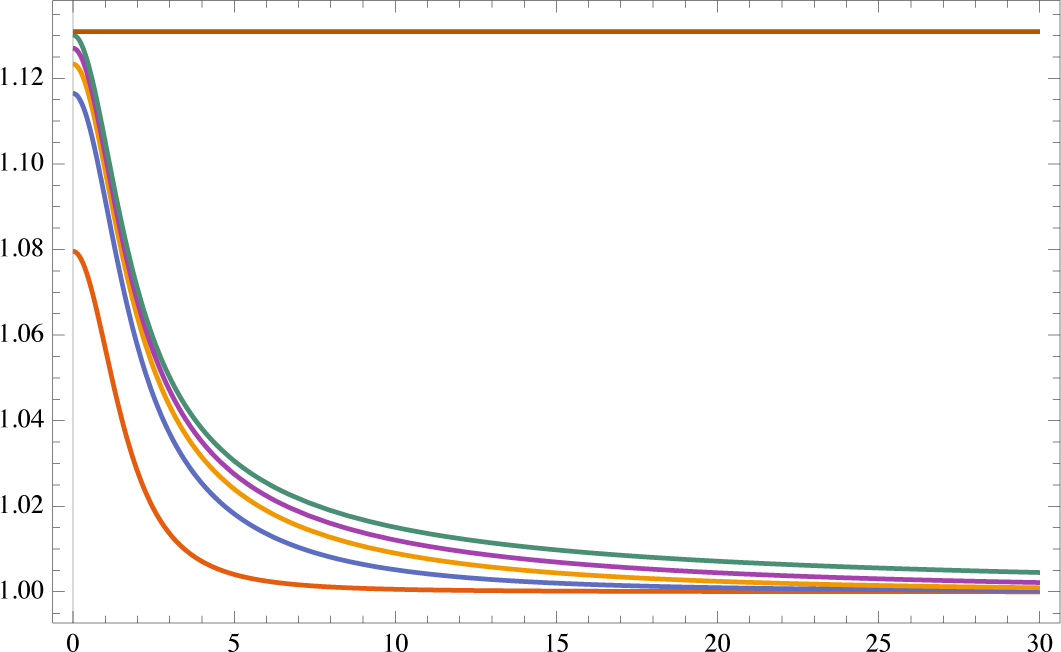
Pair correlation function *g*_*t*_(*r*) for a Cauchy kernel with parameter *b* = 1 for a process with *λ* = 0.05, *V* = 0.1 (*V/λ* = 2) computed for *t* = 1, 5, 10, 20, 100 generations. The pair correlation function increases with the number of generations but the series converges. Horizontal line is *y* = *g*_∞_(0) = 1 + *V π/*48*b*^2^*λ*

This feature of Malécot model was pointed out by Felsenstein which on the basis of this result dismissed the model as biologically non relevant. In [8] Shimatani introduces an interesting variant of the model, by supposing that at every generation a constant fraction of the offsprings migrate and are redistributed at random on the plane. As a consequence, the degree of clustering remains finite. In this paper, we select a few kernel functions with various features to test the relevancy of our model (see Nathan review paper on kernels [12] and [13]) and we show that the initial model is capable to show finite clustering if a long distance dispersal (LDD) kernel is used, without need of introducing the at random migration hypothesis.

#### 2.1.1. Cauchy dispersal kernel

This kernel is used for seed dispersal studies [15, 12, 19]. It can be obtained as a continuous mixture of Gaussian kernels with variance parameters distributed as the inverse of a Gamma distribution, see [14]. It is defined by

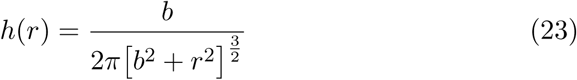

where *b* is a scale parameter. Contrarily to the Gaussian kernel, which has a finite average cluster radius given by

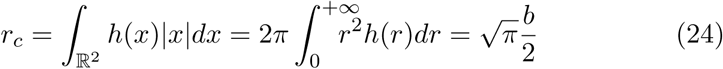

for a Cauchy kernel the above integral diverges. Therefore is is well suited for representing long distance dispersal of offsprings. In the Supplementary material we compute its 2*k*-fold convolution to be

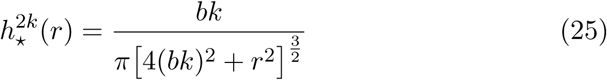

Since for *r* ≥ 0

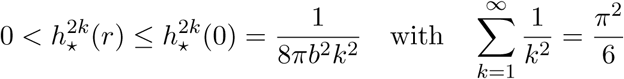

the series of functions 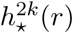 is uniformly convergent by Weierstrass’ criterion. Even if a closed form for the sum of the series is not available, we can compute the value of *g*_*t*_(*r*) in (19) for *r* = 0 and *r* → +∞ (with *U* = 1)

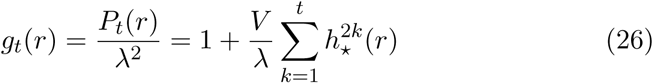

hence

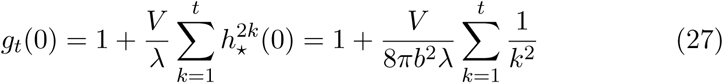

We remark that, even if the clustering is finite for a Cauchy kernel, we see that the clustering a *r* = 0 *increases* with the number of generations up to a certain limit value. In the limit *t* → ∞ we have *g*_∞_(0) = 1 + *V π/*48*b*^2^*λ*. Moreover since 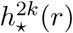 is vanishing for *r* → ∞ we have lim_*r*→+∞_ *g*_*t*_(*r*) = 1.

#### 2.1.2. Exponential kernel

It is used as a reference against more fat-tailed kernels. When used to describe pollen or seedlings dispersion it outperforms the Gaussian kernel (which is the kernel used in mechanistic diffusion or random walk models). It is defined by ([12, 13])

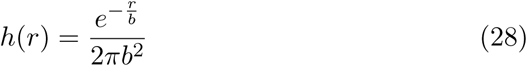

where *b* is a scale parameter. The average cluster radius is *r*_*c*_ = 2*b* and its 2*k*-fold convolution (see Supplementary material) is

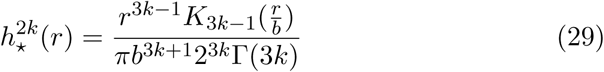

where *K*_*n*_(*r*) is the modified Bessel function of the second kind. To study qualitatively the convergence of the series of functions we adopt the asymptotic approximation *a*) for *r* → +∞ and *b*) for *r* → 0 below (see [18] Ch. 9)

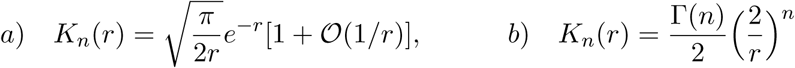

so that for large *r*

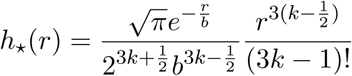

and the series of function obtained neglecting *𝒪*(1*/r*) terms converges and has closed form

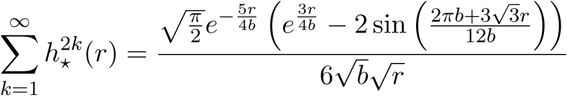

Hence for large *r*

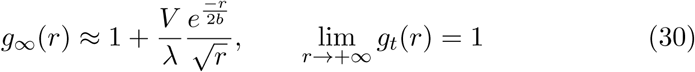

For *r* → 0 we have

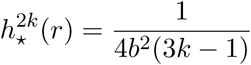

therefore

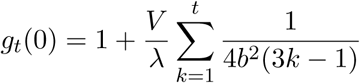

and the pair correlation function at *r* = 0 diverges for *t* → ∞ contrarily to the Cauchy kernel.

## 3. Marked point processes and spatial genetic applications

In this section we consider a replicate marked point process (*x, m*(*x*)) where the qualitative mark *m*(*x*) corresponds to the genotype of the individual at *x* and we compute the pair correlation function between individuals of given genotype. This model is related to the one in [9]. We suppose that the parent population is the sum of individuals having *m* different genotypes. For example, in the case of a single locus having *l* possible alleles, there are *m* = *l* different genotypes for an haploid individual, while there are *m* = *l*^2^ genotypes for a diploid individual. Let *λ*_*i*_ be the intensity of genotype *i* in the population, *λ* = ∑_*i*_ *λ*_*i*_ and *q*_*i*_ = *λ*_*i*_*/λ* be the frequency of genotype *i*. As before, we state that

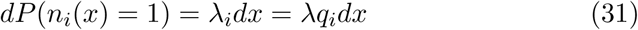

A parent of genotype *i* may have a daughter of genotype *j* with conditional probability *P* (*j|i*) = *Q*_*ij*_, where ∑_*j*_ *Q*_*ij*_ = 1, so that the probability of a *i* → *j* filiation is *Q*_*ij*_*q*_*i*_ and the frequency of genotype *j* in the first offspring generation is

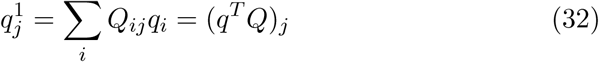

where the superscript *T* denotes transposition. If we stipulate that our reproductive mechanism is described by the time-independent Markov chain *Q* and that the genotype frequencies *q* are invariant from one generation to another, that is

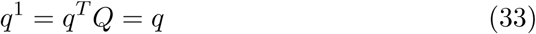

then the population is at the Hardy-Weinberg equilibrium. Moreover we assume that the dispersal mechanism described by the dispersal kernel *h* is independent of the genotype *i*. We want to compute the probability of finding two daughters of genotype *i* and *j* at distance *r* = *x*_2_ – *x*_1_. As before,

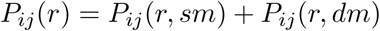

For the (*sm*) case, let *x* be the mother location. The sought probability *P*_*ij*_(*r, sm*) is the product of the probability *M*_*ij*_(*sm*) of obtaining a couple of daughters of genotype *i, j* from a mother of arbitrary genotype

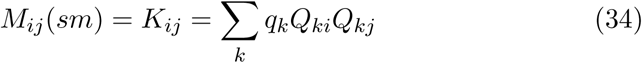

times the probability *P* (*r, sm*) in (10) since the dispersal probability kernel *h* is independent of the genotype. Hence

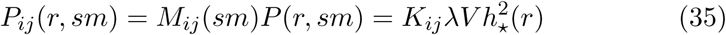

For the (*dm*) case, let *x, y* be the parent locations. We have that

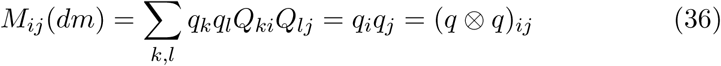

due to the stationarity of the chain (33) (Hardy-Weinberg equilibrium hypothesis). Moreover, see (12)

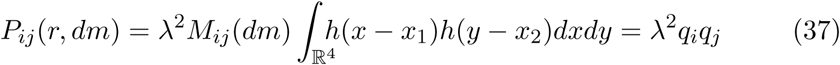

Note that in both (*sm*) and (*dm*) cases, *P*_*ij*_ = *P*_*ji*_. Summing up the above results, for the first offspring generation we have

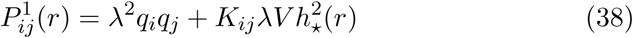

while for the subsequent generations, in the (*sm*) case, since the genotype frequency is invariant, the computation gives the same result of (10)

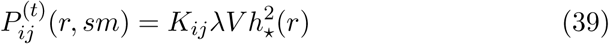

For the (*dm*) case at the *t* generation, we have

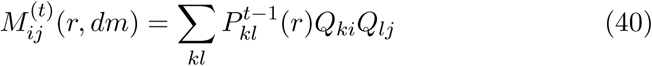

and

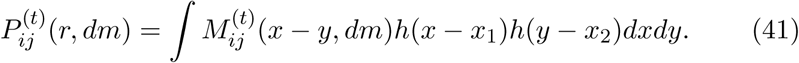

Let *Q*^*k*^ = *Q* ×*… Q* be the *k*-times product of *Q*. It is easy to see that 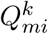 gives the probability that an ancestor of genotype *m* has a genotype *i* daughter *k* generations later. We thus denote with

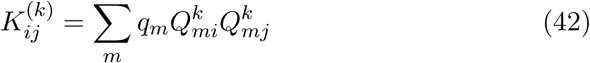

the probability of obtaining a couple of daughters of genotype *i, j* having a common ancestor *k* generations before. Let us do the computation for the *t* = 2 generation:

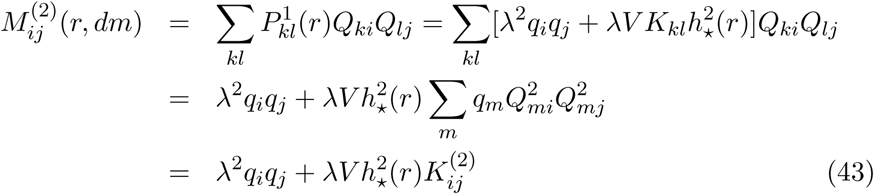

and

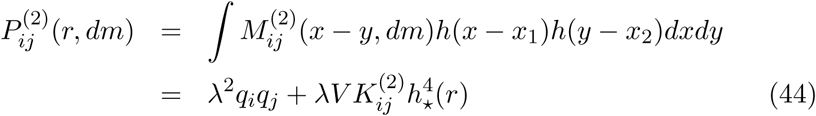

We recognize in the first term the probability of the event (*dm*)^2^ and in the second of (*sm/dm*). Therefore, by iterating the computation above, we get

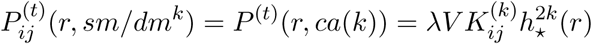

and

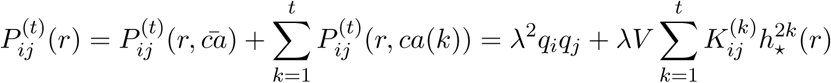

The associated pair correlation function between *i, j* genotypes is

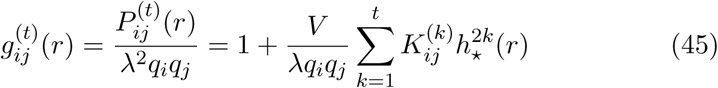

### 3.1 Ergodic case

We have already assumed that the genotype frequencies *q* are the stationary (Hardy-Weinberg) equilibrium distribution of the Markov chain *Q*. If we suppose that (*Q*^*v*^)_*ij*_ *>* 0 for some integer *v*, then by the Ergodic Theorem for Markov Chains (see e.g. [16]), we have that

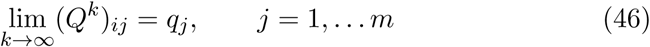

hence 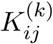 in (42) becomes 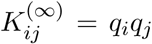 in the infinite generations limit, and all the pair correlation functions tend to

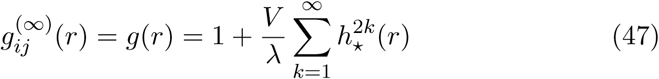

### 3.2. Distance decay of genetic similarity

Isolation by distance is the phenomenon of increasing genetic diversity between two localized subpopulations that are geographically distant or separated by a barrier. If we consider species composition in a given area instead of genetic diversity, the same concept is termed species turnover or *β*-diversity. Many drivers of diversity such as distance, ecological niches or barriers may act at the same time. Point process models are particularly effective to describe the role of clustering on the decay of similarity. A a commonly used statistical estimator is Sørensen similarity index [22] between two regions *A* and *B*

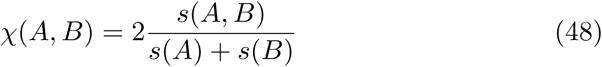

where *s*(*A, B*) is the number of species present in *A* and *B* while *s*(*A*) is the number of species in *A*. This formula has been rewritten in the point process framework in [19] in case *A* and *B* are infinitesimal regions *dx* at distance *r* as

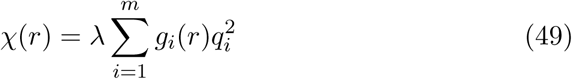

where *m* is the species number, *g*_*i*_(*r*) is the pair correlation function of species *i* and *q*_*i*_ is the frequency of species *i*. This formula can be used to measure decay of genetic similarity with distance if *m* is the number of genotypes and *g*_*i*_(*r*) is given by 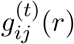 in (45) for *i* = *j*. Hence

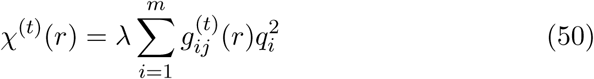

or equivalently

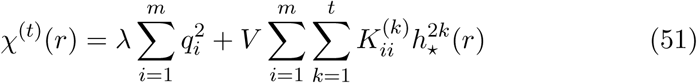

In the infinite distance limit *r* → ∞ the second term in (51) vanishes and the above *χ*^(*t*)^(*r*) tends to 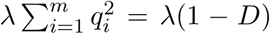 where 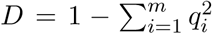 is the Gini-Simpson index. It represents the similarity at a distance where the clustering has no effect and the spatial distribution of individual can be considered completely random. Interestingly, in the ergodic case and in the infinite generations limit 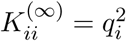 and the above *χ*^(*t*)^(*r*) tends to

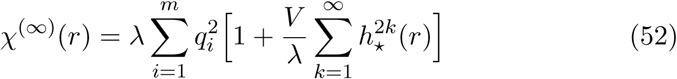

showing that there is a residual contribution to the similarity due to the clustering.

### 3.3. Spatial autocorrelation of genotypes

To make the paper self-contained, we briefly recall the notion of Moran’ index of spatial correlation [17]. We consider a lattice of *n* sites and let *a*_*i*_ be the value of a numerical variable at site *i*. The average and variance of *a* are

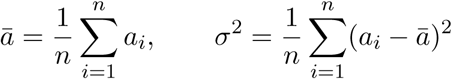

Let *W* = *w*_*ij*_, *i, j* = 1,*…, n* be a matrix with non negative entries giving a weight *w*_*ij*_ to every couple of sites, *|W |* = ∑_*ij*_ *w*_*ij*_ the sum of weights and

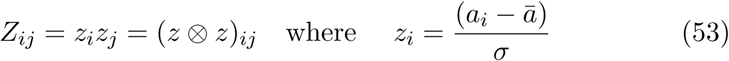

the score matrix. Moran index is

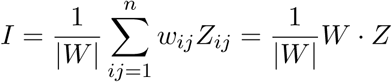

If we consider only sites with distance(*i, j*) = *|i–j|* = *r* we obtain the spatial auto-correlation of variable *a* at distance *r*

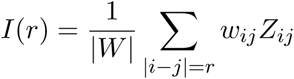

Moran index has been rewritten in the language of point processes by Shimatani in [9] by substituting i) regular lattice sites *i* with a distribution *x* of points in the plane (spatial point process), ii) the variable *a* with the quantitative mark *m*(*x*) of point *x* and iii) *w*_*ij*_ with *P* (*x, y*). To establish a direct link with the genetic model exposed in the section above, we suppose that to every genotype *i* = 1,*…, m* there corresponds a number *a*_*i*_, which can be considered a *quantitative* mark *a*(*m*(*x*)). Moreover we consider the average and variance of *a* with respect to the frequency vector *q*

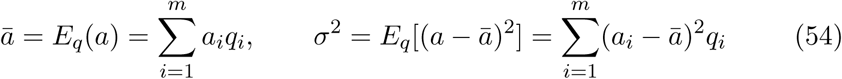

and *w*_*ij*_ = *P*_*ij*_(*r*) = *λ*_*i*_*λ*_*j*_*g*_*ij*_(*r*) given by (45) with total weight

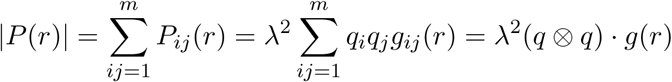

If we want to compute the value of the index *I* at generation *t*, we have to plug into the above formula 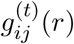 in (45). Hence, the point process version of Moran Index (see [9], [2]) is

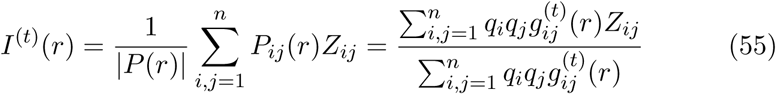

While the score matrix depends only on the average and variance of *a* with respect to the equilibrium frequencies *q*, the pair correlation matrix 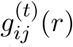 depends on distance and generations. For *r* going to ∞ the correlation between individuals vanishes (*P*_*ij*_ = *q*_*i*_*q*_*j*_, *g* = 1) hence Moran index tends to (∑_*i,j*_ *q*_*i*_*q*_*j*_ = 1)

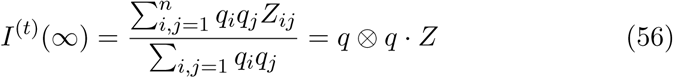

On the other hand, if the ergodic hypothesis (46) holds, then 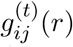 tends to *g*(*r*) in (47) for *t* → +∞ hence again Moran index become independent of the clustering of the process

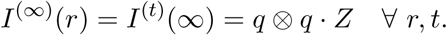

It is instructive to examine in detail the form of the correlation index *I*^(*t*)^(*r*) for finite distance *r*. By plugging into (55) the form of the pair correlation function 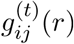 in (45), we get, after simple manipulations and using the fact that the power *Q*^*k*^ of a Markov matrix *Q* is Markov for all *k*

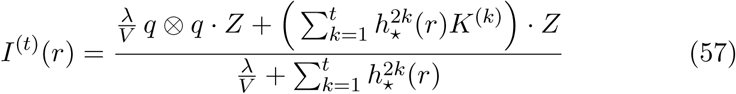

The above limits can be recovered from the last formula since 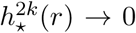 for *r* → ∞ and *K*^(*k*)^ → *q* ⊗ *q* for *k* → ∞. By rewriting (57) as

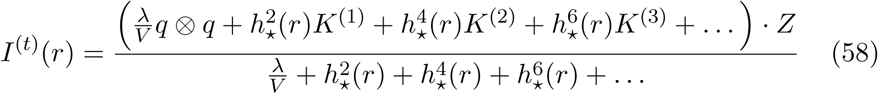

we see that the spatial correlation at very small distances *r* ≈ 0 depends on the ratio *λ/V* and on the value of the dispersal kernel 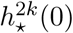. For a Cauchy kernel we have

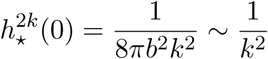

while for an exponential kernel we have

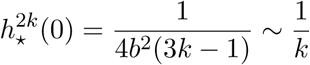

At very high spatial density of individuals *λ* the contribution of past generations is weak (see example) and the value of the index tends to the value of the uncorrelated case *q* ⊗ *q · Z*. At very low spatial density of individuals the effect of past generations can be strong depending on the type of kernel. For a Cauchy kernel representative of a situation where offsprings migrate at very large distances from the parents, only the first few generations affect the value of the index, while for an exponential kernel past generations have a greater weight. See the following example based on a similar one in [9]

### 3.4. Example

Let us consider a marked point process where the qualitative mark at a single locus of a diploid individual is defined by the following genotypes: 1 = *AA*, 2 = *A*∗ and 3 = ★★. We transform the qualitative mark into a quantitative one by the following one-to-one map *a*(1) = 1, *a*(2) = 1*/*2, *a*(3) = 0. Given a value *a ∈* (0, 1) representing the frequency of allele *A* in the population, we suppose that the genotype frequencies are

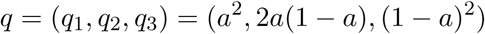

The average and variance of *a*, defined by (54) are

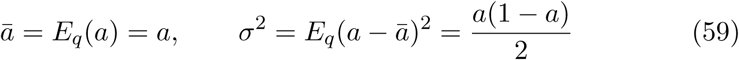

Let the mutation probabilities be given by the following Markov transition matrix

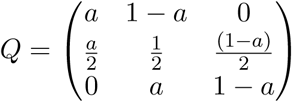

One can check that *q* is the equilibrium distribution for *Q* i.e. *q* = *q*^*T*^ *Q* see (33), hence, genotypes frequencies and their variance are invariant from one generation to the other. Moreover, since 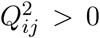, the hypotheses of the Ergodic Theorem for Markov chains (46) are fulfilled and it holds that 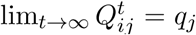. We may now compute the score matrix *Z* using (53) and the matrices *K*^(*k*)^ using (42). We get

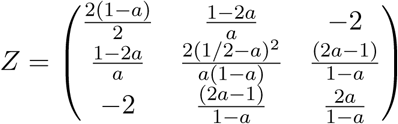

A straightforward computation shows that for this system *q* ⊗ *q · Z* = 0 meaning that in absence of clustering *P*_*ij*_(*r*) = *q*_*i*_*q*_*j*_ and the genotypes show no spatial correlation. By computing *Q*^*k*^ and *K*^(*k*)^ in (42) for this system we find that *K*^(*k*)^ *· Z* = 1*/*4^*k*^ (independent of *a*) hence Moran index in (57) become in this case

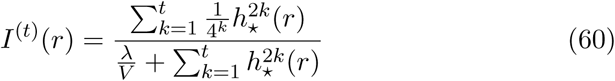

### 3.5. Comparison of Moran index for different kernels

In this Section we compare the plot of Moran Index for Gaussian, exponential and Cauchy kernels. We set the kernel parameters in order to have the same average cluster radius, when possible. For a Gaussian kernel the average cluster radius is 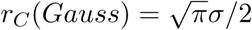, while for an exponential kernel it is *r*_*C*_(*Exp*) = 2*b*. We therefore set 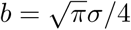. For a Cauchy kernel *r*_*C*_ (*Cauchy*) = +∞, therefore we set *b* = 1 and *b* = 4 for comparison.

*Gaussian kernel* : We see (Figure 4) that the genetic clustering decreases rapidly with the number of generations in a range 0 *≤ r ≤* 6*r*_*C*_ (*Gauss*) and it vanishes for couple of individuals at larger interdistance.

**Figure 3:**
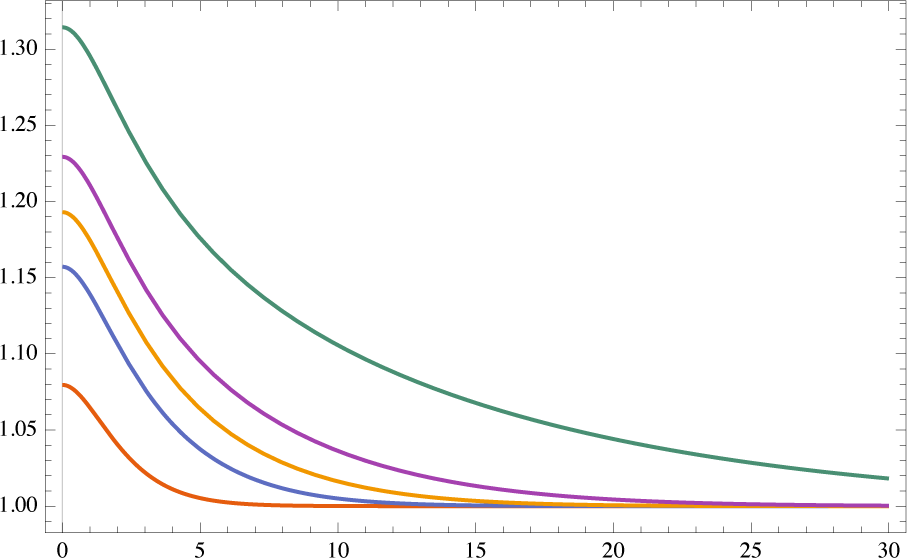
Pair correlation function *g*_*t*_(*r*) for an Exponential kernel with parameter *b* = 1 for a process with *λ* = 0.05, *V* = 0.1 (*V/λ* = 2) computed for *t* = 1, 5, 10, 20, 100 generations. The pair correlation function increases with the number of generations, diverges for *r* → 0 but converges for large *r*

**Figure 4:**
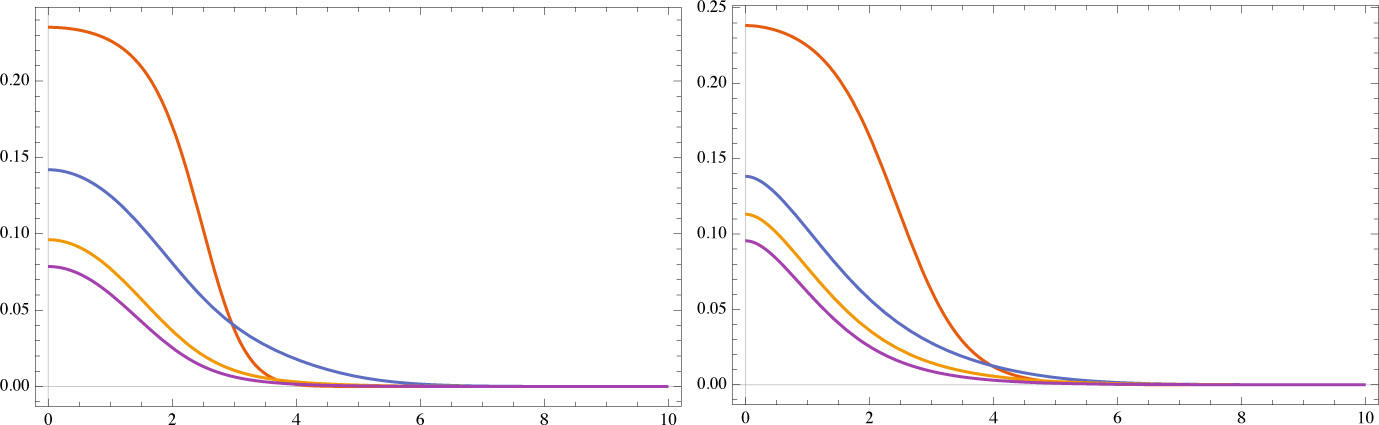
Left: Moran Index for a Gaussian kernel with parameter *σ* = 1 for a process with *λ/V* = 0.01 and *t* = 1, 5, 10, 20. Right: Moran Index for an Exponential kernel with parameters 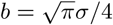, *σ* = 1 for a process with *λ/V* = 0.01 and *t* = 1, 5, 10, 20. Curves are decreasing with the number of generations *t*. For both kernels there is a steep decay in the genetic clustering for *r >* 5*r*_*C*_

*Exponential kernel* (Figure 4): the behavior is similar to the Gaussian kernel case, but the genetic clustering at *r* ≈ 0 is higher and the index shows little variation for *t >* 10, hence the Moran index can be considered close to the limit value for *t* ≥ 20

*Cauchy kernel* (Figure 5): the curves have a longer tail, hence the genetic relatedness between individuals is nonzero even for larger interdistances. Moreover, the index is clearly close to its limit value for *t* ≥ 20.

**Figure 5:**
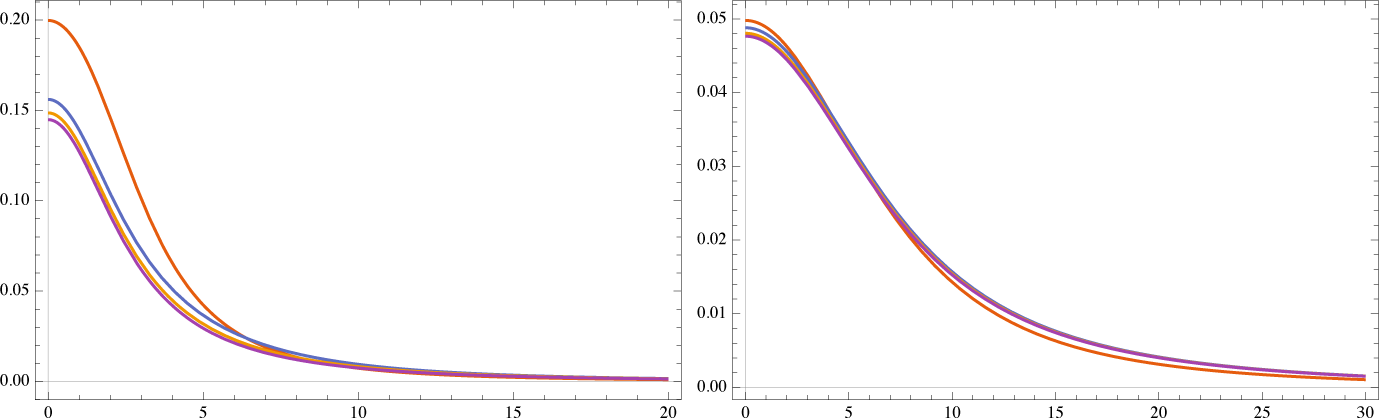
Moran Index for a Cauchy kernel with parameter *b* = 1 (left) and *b* = 4 (right) for a process with *λ/V* = 0.01 and *t* = 1, 5, 10, 20. Curves are decreasing with *t*. The fat tail in the curves indicates that there is slow decay of spatial genetic correlation with the interdistance

As a general feature, spatial correlation between individuals at certain interdistance *r* as measured by the pair correlation function *g*_*t*_(*r*) *increases* with the number of generations (see e.g. Figure 3) while the genetic relatedness as measured by Moran spatial auto correlation index *I*^*t*^(*r*) or by the average kinship coefficient *G*_*t*_(*r*) *decreases* with the number of generation. This is in accord with the results in [9], and at contrast with previous theoretical studies on lattice models (see [9]). However, at a finer scale the behaviour of these statistical descriptors do depend on the form of the dispersal kernel function as shown in this work. Hence, the joint use of pattern and genetic descriptor allows a better design and validation of spatially explicit models to describe population genetics and spatial ecology models (sese e.g. [11]).

### 3.6. Kinship coefficient

A commonly used measure of genetic relatedness is the so called kinship coefficient *ϕ* between two arbitrarily chosen individuals: it gives the probability that two randomly chosen alleles from the same autosomal locus are identical by descent, i.e they are identical and they come from the same ancestor, see [21]. The kinship coefficient can be computed given the pedigree. *ϕ* is 1*/*4 for a couple of full siblings or for a couple parent-offspring, it is 1*/*8 for a couple of half-sib (only the mother is in common). In our model we have only maternal half-sib pairs. *ϕ* is zero in the (*dm*) case because the two individuals are not genetically related, it is 1*/*8 in the (*sm*) case and it is

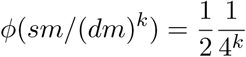

for two individuals having a common ancestor *k* generations ago. In [8] Shimatani computes the average kinship coefficient for a couple of individuals at distance *r* along *t* generations as

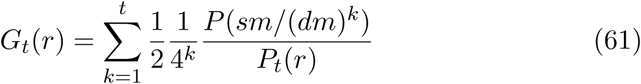

By substituting (15) and (16) in the above formula we get

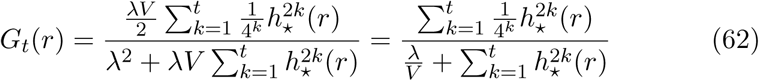

Note that *G*_*t*_(*r*) in the above formula coincides with Moran Index *I*^(*t*)^(*r*) in (60). Therefore the above plots in Figures 4 and Figure 5 can be used to compare the average kinship coefficient for different kernels type.

### 3.7. Founder effect

The founder effect [23, 24] is the loss of genetic variation that occurs when a new population is established by a very small number of individuals segregated from the original larger population. Being small in size the new population is not a representative sample of the ancestral population, moreover it will undergo considerable genetic differentiation due to the accumulation of inbreeding caused by the isolation.

We show how our model can be tuned to model the founder effect. We mimick the segregation effect by using a short range dispersal kernel and the effect of inbreeding by considering high population density (intensity). Looking at short interdistance (small size community), we want to see if there is still a high value of Moran index after a few generations. From Figure 4 we already know that genetic clustering (spatial correlation) will eventually decrease when the number of generations *t* become large.

On the one hand, from Figure 6 we see that for Gaussian and Exponential kernel, which are short range dispersal kernel, Moran index remains considerably high also when population density is increased 100 times in a interdistance range between 0 and 5 times the average cluster radius. On the other hand, (see Figure 7) using the Cauchy long dispersal range kernel there is a strong decrease of genetic clustering in the same interdistance range at higher population density. The large dispersal distance property of this kernel prevents the segregation effect and enhance the genetic mixing therefore it prevents the founder effect to display.

**Figure 6:**
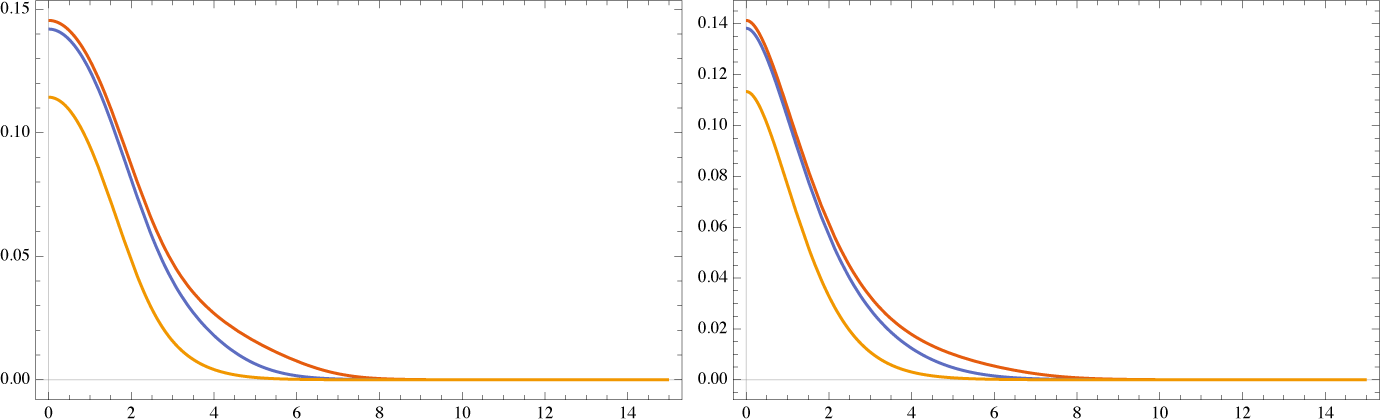
Moran index for different kernels and increasing intensity. Left: Gaussian kernel with *σ* = 1 and intensity *λ*_0_ = 10^*-*3^, *λ*_1_ = 10^*-*2^, *λ*_2_ = 10^*-*1^, Right: Exponential kernel with 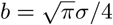 and intensity *λ*_0_, *λ*_1_, *λ*_2_. Curves are decreasing with *λ*

**Figure 7:**
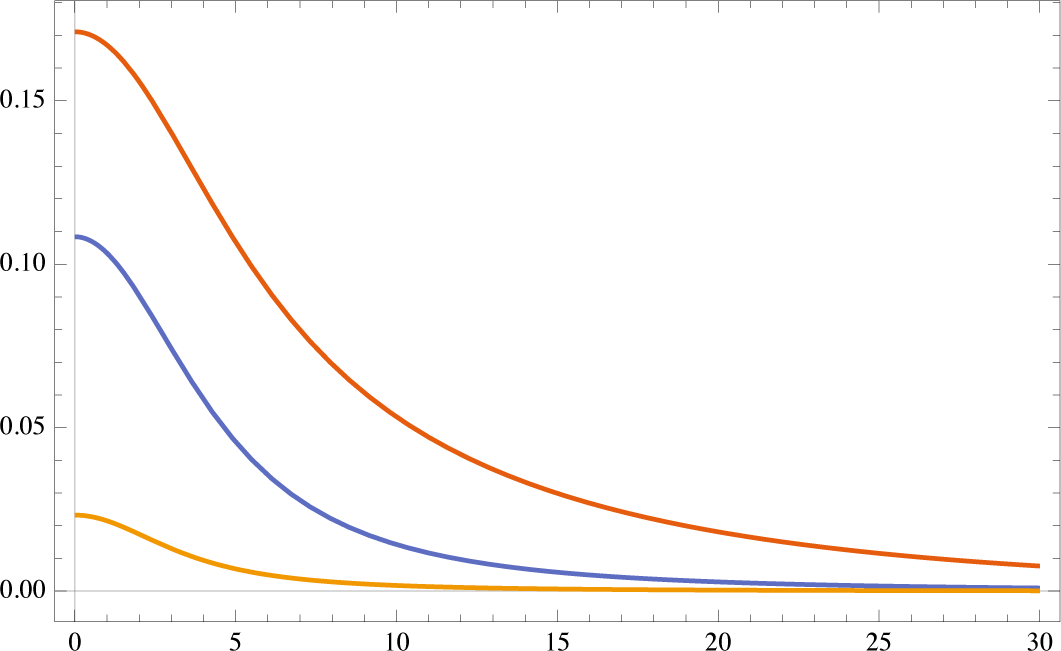
Moran Index for a Cauchy kernel with *b* = 2 and intensity *λ*_0_ = 10^*-*3^, *λ*_1_ = 10^*-*2^, *λ*_2_ = 10^*-*1^. The curves is decreasing with *λ* and show a strong decay in the genetic clustering at short interdistance.

In Figure 8 we show the flexibility of the model with respect to the dispersal kernel parameters: using a Gaussian and exponential kernel with an average cluster radius 6 times higher of the one used in Figure 6 and a scale parameter *b* = 1 for Cauchy kernel half the value of the one used in Figure 7 we obtain a comparable behavior for the three kernels.

**Figure 8:**
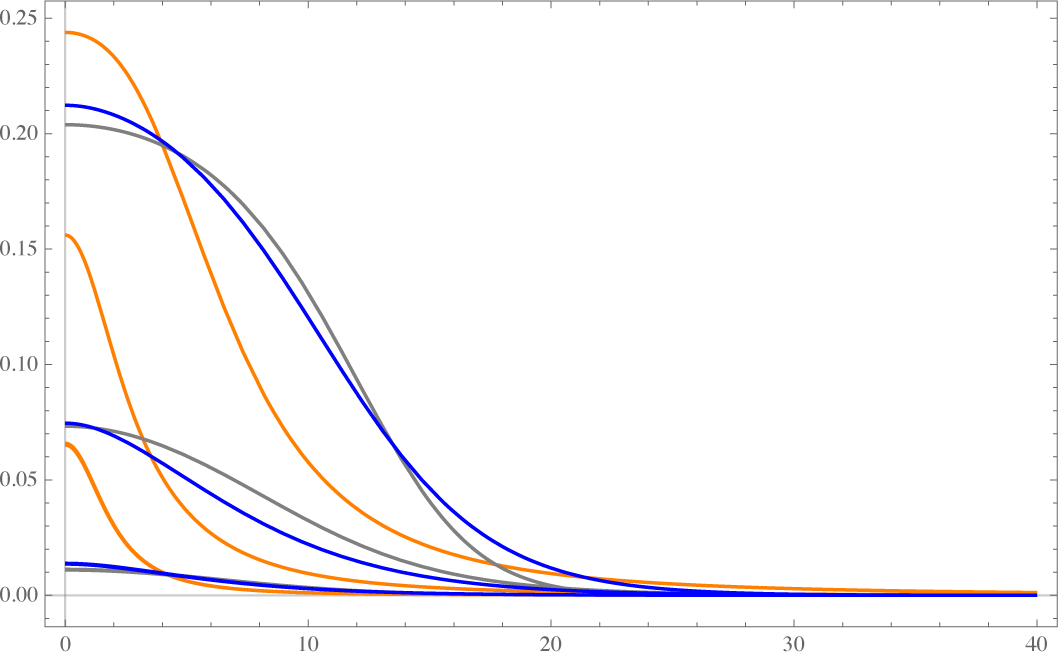
Cumulative illustration of the founder effect. Moran Index computed for a Cauchy kernel (*b* = 1, orange), Gauss kernel (*σ* = 6, gray) and Exponential kernel (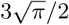, blue). Intensity is increased with the number of generations: *λ/V* = 0.001 for *t* = 1, *λ/V* = 0.01 for *t* = 5, *λ/V* = 0.1 for *t* = 10 and *t* = 20. For all kernels curves are decreasing with *t* and *λ*. Using for the Gaussian and exponential kernel an average cluster radius 6 times the one used in Figure 6, and a scale parameter *b* = 1 half the value of the one used in Figure 7 we obtain a comparable behavior for the three kernels.

## 4. Conclusions

The starting point for this work is the replicated Neyman-Scott point process model in [8]. The main weak point of the model in [8] and of previous papers is the use of a short dispersal range Gaussian kernel. In this way the clustering diverges with the number of generations. Here we have computed the pair correlation function of the replicated point process for a general dispersal kernel and given sufficient conditions for the convergence of the clustering. Clustering is finite for leptokurtic (fat tailed) dispersal kernels, which in many studies outperform the Gaussian or exponential kernels at fitting seed dispersal data. The result in this paper has an immediate application e.g. to the decay of similarity model introduced in [19], which is based on the pair correlation functions of the various species.

Replicated marked point process models considered in [8] are important because they allow to discuss simultaneously spatial and genetic variability patterns. In this way spatial distribution and population genetic information can be used jointly to pinpoint the model and select the form of the dispersal kernel, a task which is not easily performed for fat tailed kernels if only spatial data are available.

In Section 3 we have discussed a simple model of replicated point process of genotyped individuals where the genotype frequencies at equilibrium are expressed by a discrete time, finite state Markov chain. We have computed pair correlation function for genotyped individuals (called mark correlation function in [2]). We have discussed with examples the role of the dispersal kernel in this model for describing classical statistical estimators as Moran Index of spatial correlation of genotypes or the average kinship coefficient. The possibility of using different kernels gives a great flexibility to the model and allows to discuss classical notions as the founder effect. In particular, the divergence of the clustering for short range kernels in the infinite generations limit seems not to be a serious limitation when modeling actual systems of community ecology. We think that the techniques discussed here have a great potential and a wide range of applicability in population genetic and community ecology studies. A next step would be to relax the assumption of constant intensity (density of individuals) at every generation, in order to include in the model important effect such as bottlenecks or extinctions.

## Competing Interests

The author declares no competing financial interests.

## 5. Supplementary Material

### 5.1. Replicated Neyman Scott process with a radial dispersal kernel

The two-dimensional stationary and isotropic Neyman-Scott process is defined by conditions i) -iii) below:

i) parents are distributed over the plane following a homogeneous Poisson process with density *λ*
ii) each parent produces *u* daughters where *u* is a random variable with known distribution, Let *U* = 𝔼 (*u*) be the expectation and *V* the variance
iii) daughters are dispersed around the mother according to a radial dispersal kernel *h*(*r*) with 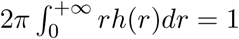

Parents are removed from the plane at each generation produced by steps ii) and iii). The main object to describe a clustered distributions of individuals in the plane is the product density

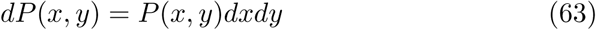

which is the probability of finding two individuals in the infinitesimal areas *dx* and *dy* located by the vectors *x* = *x* – 0 and *y* = *y* – 0 in the plane, where 0 is an arbitrarily fixed origin in the plane. As usual *dP* (*x*) = *λdx* is the probability of finding an individual located at *x*. We denote with *x* the point of the plane and with *x -* 0 the vector from the origin to the point if there is risk of confusing them. The pair correlation function is defined as

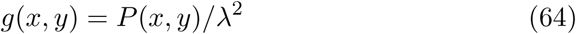

where *λ* is the density of individuals. Both can be computed given the dispersal kernel function which gives the probability that a daughter of a mother at *x* can be found (lands) at *u*

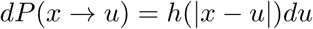

where we have assumed that *h*(*|x – y|*) is a radial function, hence *h*(*r*) = *h*(*-r*).

Given two points *x*_1_ and *x*_2_ we want to compute the probability *P*_1_(*x*_1_, *x*_2_) of finding two first generation daughters at these points. We have to consider two cases: dm) daughters have two different mothers located at *x* and *y* respectively and sm) daughters have the same mother located at *x*. Introduce for the case dm) the vectors

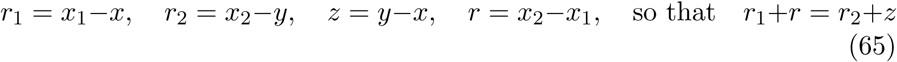

and for the case sm)

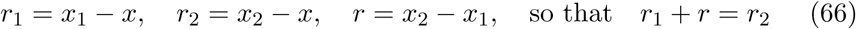

The sought probability *P* (*x*_1_, *x*_2_) can be seen as the sum of two mutually incompatible and exhaustive alternatives: *dm* (different mother) and *sm* (same mother) thus

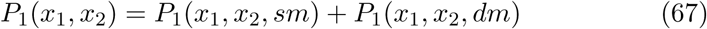

We have

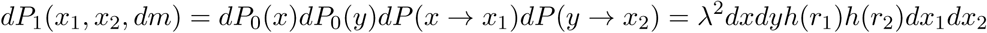

Since on the average a mother produces *U* daughters and the two mothers can be located everywhere in the plane we have to multiply by *U*^2^ and integrate with respect to the area measures *dx* and *dy*. Set for simplicity sake

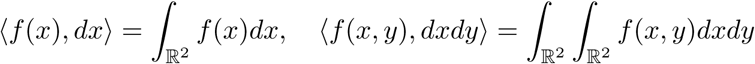

and suppose that Fubini Theorem holds everywhere so that

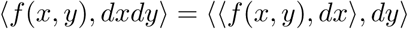

Moreover recall that for a dispersal kernel 〈*h*(*r*), *dr*〉 = 1. Using the change of variables in (65) *x*_1_ = *r*_1_ + *x* and *x*_2_ = *r*_2_ + *y* we can integrate with respect to *dr*_1_ = −*dx*_1_ and *dr*_2_ = *-dx*_2_ so that in the *dm* case we have

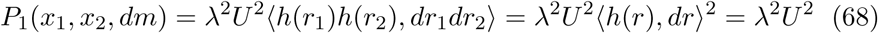

independent of *x*_1_, *x*_2_. For the *sm* case we have

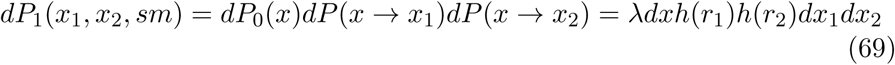

We have to multiply by 𝔼(*u*(*u* – 1)) = *WU*^2^ where *W* = (*V* + *U*^2^ – *U*)*/U*^2^ and use the change of variables *x*_1_ = *x* + *r*_1_ in (66) to integrate in −*dr*_1_ hence

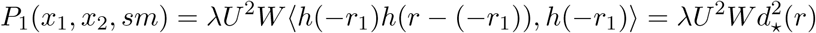

where we have introduced the convolution of two functions on the plane *f* (*x*) and *g*(*x*)

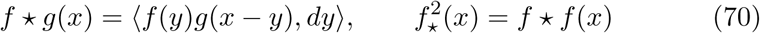

Note that the convolution of two radial functions is a radial function. Collecting the previous results we have

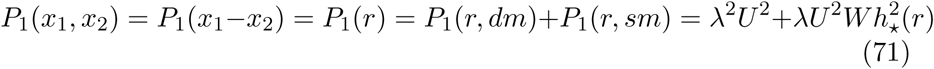

Note also that since *P*_1_(*r*) = *P*_1_(*r, dm*) + *P*_1_(*r, sm*) then

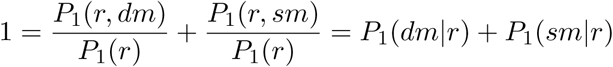

Now we consider the generation *n* = 2. Note that at each generation the kernel terms like e.g *dP* (*x* → *x*_1_) do not change, the terms *P*_*n*_(*x*)*dx* have to be updated with the new parent density *λU*^*n*^ and the terms *P*_*n*_(*x, y*)*dxdy* are the result of the *n* – 1 step calculation. Hence

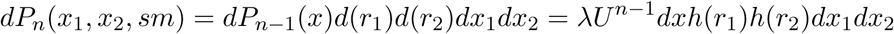

and the same computation in (69) gives the result

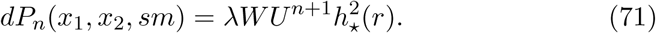

For the dm) case, we use the result (68). Hence

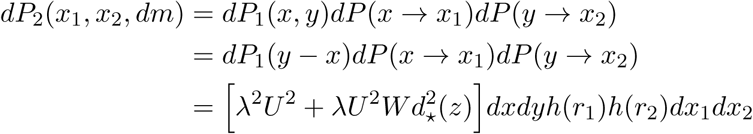

we have to multiply by *U*^2^ and integrate in *dx* and *dy*. The first term is as before

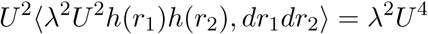

while for the second since *z* = *r*_1_ + *r – r*_2_ and *dx* = −*dr*_1_, *dy* = −*dr*_2_

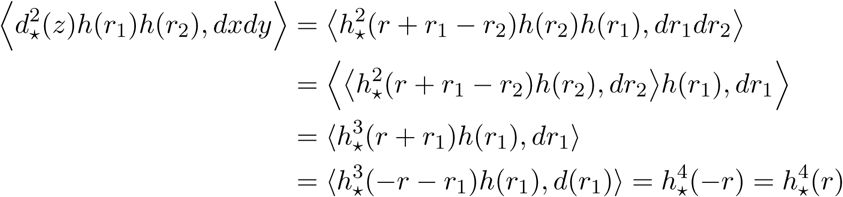

hence

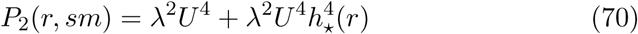

and

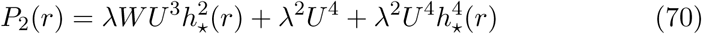

In the general case, we have the recursive formula with *z* = *r*_1_ + *r – r*_2_ 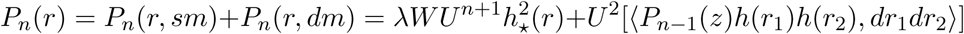 One can check that

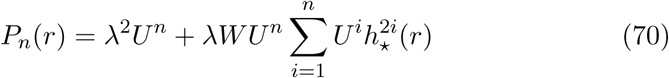

and, using the *n* generation density *λU*^*n*^, that

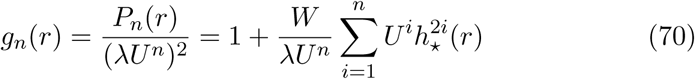

These formulae are the generalization to an arbitrary radial kernel of the results obtained by Shimatani in [8], formulae (9) and (10), for a Gaussian kernel.

### 5.2 Convolution of 2-dimensional radial functions

We briefly report here the theory necessary for computing the convolution function *f* given the dispersal kernel *h* [25]. Given a function *f* (*x*) on the plane, *x ∈* ℝ^2^, let us denote with *F* (*f*) = *f* its Fourier transform. We recall also the Convolution Theorem:

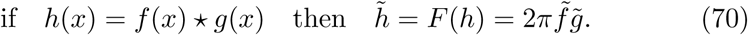

For radial functions *f* (*r*) = *f* (*|x|*) the Fourier transform can be computed^1^ using the Bessel function *J*_0_(*z*)

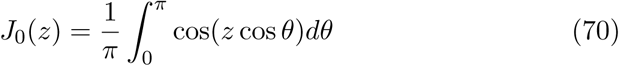

as

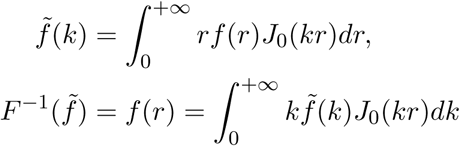

which shows that the Fourier Transform of a radial function is a radial function and vice versa.

We need to compute *h* ★ *h* where *h* is a radial function. Since

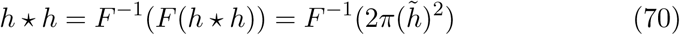

we have that *h* ★ *h* is a *radial* function that can be computed as

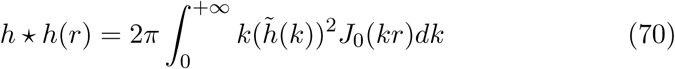

where

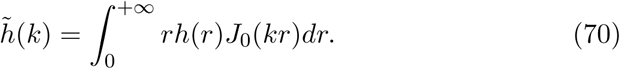

In the general case, writing 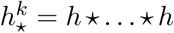 to denote the *k* -fold convolution product, we have

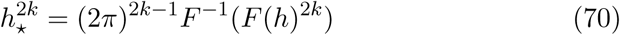

and

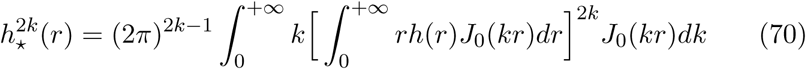

#### 5.2.1. Cauchy kernel

It is defined by

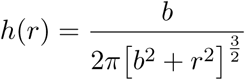

where *b* is a scale parameter. Using the above formula (5.2) we compute its 2*k*-fold convolution

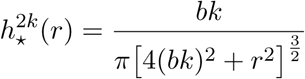

#### 5.2.2. Exponential kernel

It is defined by

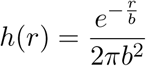

where *b* is a scale parameter. The average cluster radius is *r*_*c*_ = 2*b* and its 2*k*-fold convolution is

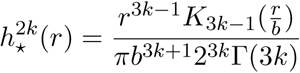

where *K*_*n*_(*r*) is the modified Bessel function of the second kind.

## Footnotes

1 Note that the Fourier transform exists only if 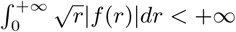.

